# Tissue-specific evolution of protein coding genes in human and mouse

**DOI:** 10.1101/011692

**Authors:** Nadezda Kryuchkova-Mostacci, Marc Robinson-Rechavi

## Abstract

Protein-coding genes evolve at different rates, and the influence of different parameters, from gene size to expression level, has been extensively studied. While in yeast gene expression level is the major causal factor of gene evolutionary rate, the situation is more complex in animals. Here we investigate these relations further, especially taking in account gene expression in different organs as well as indirect correlations between parameters. We used RNA-seq data from two large datasets, covering 22 mouse tissues and 27 human tissues. Over all tissues, evolutionary rate only correlates weakly with levels and breadth of expression. The strongest explanatory factors of purifying selection are GC content, expression in many developmental stages, and expression in brain tissues. While the main component of evolutionary rate is purifying selection, we also find tissue-specific patterns for sites under neutral evolution and for positive selection. We observe fast evolution of genes expressed in testis, but also in other tissues, notably liver, which are explained by weak purifying selection rather than by positive selection.

## Introduction

Understanding the causes of variation in protein sequence evolutionary rates is one of the major aims of molecular evolution, and has even been called a “quest for the universals of protein evolution” (Rocha 2006). Studies in a variety of organisms have reported that protein evolutionary rates correlated with many parameters, structural and functional (Pál et al. 2006; Rocha & Danchin 2004). Most notably, expression level has been shown to be the best predictor of evolutionary rate in yeasts and bacteria: highly expressed proteins are generally more conserved (Drummond et al. 2005; Pál et al. 2001; Wall et al. 2005). In animals and plants, our understanding has been complicated by the fact that genes can have different expression levels depending on tissue or life history stage, and by correlations with multiple other factors such as recombination rate, gene length or compactness, and gene duplications (Larracuente et al. 2008; Makino et al. 2009; Yang & Gaut 2011; Liao et al. 2006; Li et al. 2007). In mammals, expression breadth has been suggested to be more important than expression level (Duret & Mouchiroud 2000; Park & Choi 2010). It has also been suggested that selection against protein misfolding is sufficient to explain covariation of gene expression and evolutionary rate across taxa, including mouse and human (Drummond & Wilke 2008). This notably explains the slower evolution of brain-expressed genes; the relation with the influence of breadth of expression is unclear. Moreover, it was shown that conserved sites and optimal codons are significantly correlated in many organisms, including mouse and human (Drummond & Wilke 2008).

These correlations of evolutionary rate to many other parameters, which are themselves correlated (e.g., gene length and GC content), poses problems to determining what is true and what is spurious correlation. To disentangle which factors could be determining evolutionary rates a solution is to use partial correlation, taking into account the relationship between gene structure and other parameters when considering the correlations with evolutionary rate (Larracuente et al. 2008; Warnefors & Kaessmann 2013).

Here we aim to disentangle aspects of protein evolutionary rate and its explanatory factors in human and mouse. We use partial correlations, taking into account not only different structural parameters, but also different aspects of gene expression (level, tissue specificity), using expression in more than 20 tissues. We also used three measures of protein evolutionary rate estimated from the branch-site model (Zhang et al. 2005): strength of negative selection (value of dN/dS on sites under negative selection); proportion of neutrally evolving sites; and evidence for positive selection. This allows us to distinguish fast evolution due to weak purifying selection from that due to positive selection.

## Materials and Methods

We used RNA-seq data for mouse from the ENCODE project (The ENCODE Project Consortium 2011) and for human from Fagerberg et al. (Fagerberg et al. 2013). For mouse, the raw reads in FASTQ format obtained from the ENCODE FTP server (ftp://hgdownload.cse.ucsc.edu/goldenPath/mm9/encodeDCC/wgEncodeCshlLongRnaSeq/) were processed with TopHat and Cufflinks (Trapnell et al. 2012), using the gene models from Ensembl version 69 (Flicek et al. 2013). For human, processed data from Fagerberg et al. (Fagerberg et al. 2013) were retrieved from the ArrayExpress database (E-MTAB-1733) (Rustici et al. 2013). 22 tissues for mouse and 27 tissues for human were analyzed, of which 16 are homologous between the two species. Processed RNA-seq expression was further treated as follows (R script in Supplementary Material): multiplied by 10^6^(to avoid values under 1, which are negative after log transformation); log_2_ transformation; and quantile normalization, replacing zero values by log_2_(1.0001).

We used either global parameters of expression: median expression, maximal expression, and specificity among all tissues; or expression in each tissue separately. Expression specificity τ was calculated as follows (Yanai et al. 2005), where *x* is the vector of expression levels over all *n* tissues for a gene:

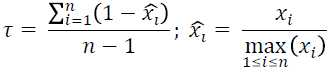

Values of expression specificity close to zero indicate that a gene is broadly expressed, and close to one that it is specific of one tissue.

Additional analysis was performed on microarray expression data from 22 human and 22 mouse tissues selected from the Bgee database (Bastian et al. 2008), as well as 8 human and 6 mouse tissues from Brawand et al. RNA-seq (Brawand et al. 2011). The corresponding results are presented in Supplementary Materials.

As measures of evolutionary rate of protein-coding genes, we used the estimates from the branch-site model (Zhang et al. 2005) as precomputed in Selectome (Moretti et al. 2014): purifying selection estimated by the dN/dS ratio ω_0_ on the proportion of codons under purifying selection (noted “Omega” in the figures), evidence for positive selection estimated by the log-likelihood ratio ΔlnL of H_1_ to H_0_ (models with or without positive selection), and the proportion of neutrally evolving sites p_1_. The evolutionary rate parameters were estimated from the Euteleostomi gene trees on the Murinae branch for mouse and the Homininae branch for human. We also present in Supplementary Materials another estimation of evolutionary rate, using the exon based MI score (Rodriguez et al. 2013; Ezkurdia et al. 2014).

For all parameters the longest coding transcript was chosen as a representative of the gene, as the evolutionary rate data were available only for this transcript. Analysis was also redone for mouse using the most expressed transcript; results are presented in Supplementary Materials. Intron number, intron length, CDS length (coding sequence length) and GC content were taken from Ensembl 69 (Flicek et al. 2013). Essentiality data were manually mapped and curated (Walid Gharib, personal communication) for human from the OMIM database (McKusick-Nathans Institute of Genetic Medicine 2014) and for mouse from the MGI database (Blake et al. 2014).

Data of expression at different developmental stages were obtained from Bgee (Bastian et al. 2008). The parameter stage number indicates the number of stages in which the gene was reported as expressed. Mouse development was divided in 10 stages: 1. Zygote 2. Cleavage 3. Blastula 4. Gastrula 5. Neurula 6. Organogenesis 7. Fetus 8. Infant 9. Adolescent 10. Adult; and human in 8 stages: 1. Cleavage 2. Blastula 3. Organogenesis 4. Fetus 5. Infant 6. Adolescent 7. Early adult 8. Late Adult.

Phyletic age and connectivity (protein-protein interactions) data were downloaded from the OGEE database (Chen et al. 2012), as ordinal data. Phyletic age stages used are: 1. Mammalia 2. Chordata 3. Metazoa 4. Fungi/Metazoa 5. Eukaryota 6. Cellular organism. Recombination rate was calculated from Cox et al. 2009 data (Cox et al. 2009).

Correlation between the different parameters was performed in two ways: simple pairwise Spearman correlation and partial Spearman correlation (results for Pearson correlation are also presented in Supplementary Materials). For partial correlation each pair of parameters were compared taking into account all other parameters: first a linear model according to all other parameters for each of the two analyzed parameters was calculated; then the Spearman correlation was calculated on the corresponding residuals. All R code is available as Supplementary Materials.

Partial correlation was used to determine the correlation between two parameters excluding dependencies from other parameters. The principle of the partial correlation can be shown on a toy example (Supplementary table S1). As example data for human height and leg length were simulated, so that either a) the length of both legs is calculated depending on height, or b) the length of the left leg is calculated from height, and the length of the right leg is calculated from the length of left leg. With simple correlation the two cases cannot be distinguished, as all three parameters correlate strongly with each other. With partial correlation we can distinguish the two cases: in case a) left leg and right leg length don’t correlate with each other if we exclude influence of the height, but in case b) we see a strong correlation between them, as expected, while right leg length no longer correlates with height. Expression, intron length, intron number, CDS length, τ, ω_0_, paralog number were log_2_ transformed before calculations. p_1_ and ΔlnL were normalized by taking the fourth root (Canal 2005; Roux et al. 2014). For parameters containing zeros a small value was added before log transformation, chosen as the minimal non zero value of the parameter (except for RNA-seq, see detailed treatment above). Altogether 9553 protein-coding genes for human and 9485 protein-coding genes for mouse were analyzed.

All the analysis was performed in R (R Core Team 2012) using Lattice (Sarcar 2008), plyr (Wickham 2011). For the representation of the data Cytoscape version 2.8.2 (Shannon et al. 2003) with library RCytoscape (Shannon et al. 2013) and Circos version 0.62-1 (Krzywinski et al. 2009) were used.

## Results

We detail here the results of Spearman partial correlation analyses (table 1); standard Spearman and Pearson, as well as partial Pearson, correlations are provided in Supplementary Materials. Spearman correlation was preferred as most of the data analyzed are not normally distributed (supplementary fig. S1), even after transformation, and to avoid a large influence of outliers. It should be noted that parameters that are expected to have strong direct relations remain strongly correlated in the Spearman partial correlation. For example the correlation between coding sequence (CDS) length and intron number, in mouse, is ρ = 0.683 for partial vs. ρ = 0.760 for simple correlation, showing that longer genes have more introns. Similarly, partial correlations still show that higher expressed genes are broadly expressed, and that specific genes have lower expression in general. Thus little relevant information is lost, while spurious correlations can be hopefully avoided.

**Table 1.**
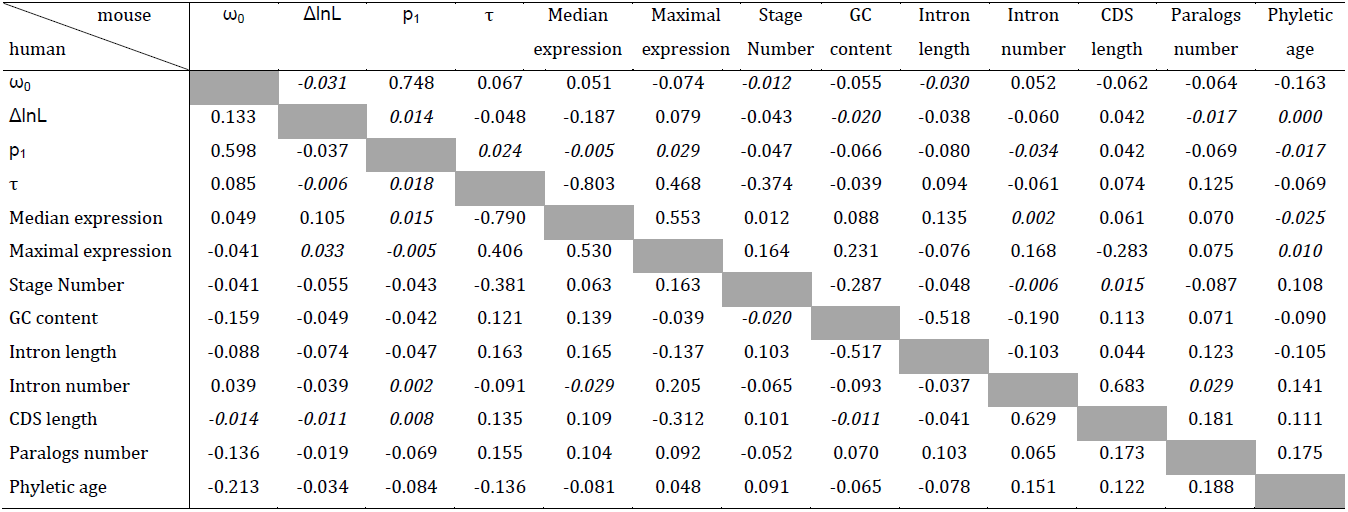
Values of partial Spearman correlations between parameters, over all tissues. Top right of table: values for mouse (corresponding to Fig 1A); bottom left of table: values for human (corresponding to Fig 1B). Not significant (p-value>0.0005) are in italics.

| mouse<br>human | $\omega_0$ | $\Delta\ln L$ | $p_1$ | $\tau$ | Median<br>expression | Maximal<br>expression | Stage<br>Number | GC<br>content | Intron<br>length | Intron<br>number | CDS<br>length | Paralogs<br>number | Phyletic<br>age |
| --- | --- | --- | --- | --- | --- | --- | --- | --- | --- | --- | --- | --- | --- |
| $\omega_0$ | | -0.031 | 0.748 | 0.067 | 0.051 | -0.074 | -0.012 | -0.055 | -0.030 | 0.052 | -0.062 | -0.064 | -0.163 |
| $\Delta\ln L$ | 0.133 | | 0.014 | -0.048 | -0.187 | 0.079 | -0.043 | -0.020 | -0.038 | -0.060 | 0.042 | -0.017 | 0.000 |
| $p_1$ | 0.598 | -0.037 | | 0.024 | -0.005 | 0.029 | -0.047 | -0.066 | -0.080 | -0.034 | 0.042 | -0.069 | -0.017 |
| $\tau$ | 0.085 | -0.006 | 0.018 | | -0.803 | 0.468 | -0.374 | -0.039 | 0.094 | -0.061 | 0.074 | 0.125 | -0.069 |
| Median expression | 0.049 | 0.105 | 0.015 | -0.790 |  | 0.553 | 0.012 | 0.088 | 0.135 | 0.002 | 0.061 | 0.070 | -0.025 |
| Maximal expression | -0.041 | 0.033 | -0.005 | 0.406 | 0.530 |  | 0.164 | 0.231 | -0.076 | 0.168 | -0.283 | 0.075 | 0.010 |
| Stage Number | -0.041 | -0.055 | -0.043 | -0.381 | 0.063 | 0.163 |  | -0.287 | -0.048 | -0.006 | 0.015 | -0.087 | 0.108 |
| GC content | -0.159 | -0.049 | -0.042 | 0.121 | 0.139 | -0.039 | -0.020 |  | -0.518 | -0.190 | 0.113 | 0.071 | -0.090 |
| Intron length | -0.088 | -0.074 | -0.047 | 0.163 | 0.165 | -0.137 | 0.103 | -0.517 |  | -0.103 | 0.044 | 0.123 | -0.105 |
| Intron number | 0.039 | -0.039 | 0.002 | -0.091 | -0.029 | 0.205 | -0.065 | -0.093 | -0.037 |  | 0.683 | 0.029 | 0.141 |
| CDS length | -0.014 | -0.011 | 0.008 | 0.135 | 0.109 | -0.312 | 0.101 | -0.011 | -0.041 | 0.629 |  | 0.181 | 0.111 |
| Paralogs number | -0.136 | -0.019 | -0.069 | 0.155 | 0.104 | 0.092 | -0.052 | 0.070 | 0.103 | 0.065 | 0.173 |  | 0.175 |
| Phyletic age | -0.213 | -0.034 | -0.084 | -0.136 | -0.081 | 0.048 | 0.091 | -0.065 | -0.078 | 0.151 | 0.122 | 0.188 |  |

### Evolutionary rate: global influences on selection

Evolutionary rate is represented by three parameters in this study, taken from the branch-site model (see Methods): ω_0_ = dN/dS, measures the intensity of purifying selection on the subset of sites determined to be under purifying selection; p_1_ is the proportion of neutrally evolving sites; and ΔlnL measures the strength of evidence for positive selection.

In both mouse and human, none of the aspects of gene expression yields a strong partial correlation to any feature of evolutionary rate (table 1; fig. 1). There is a weak correlation of ω_0_ to expression specificity τ in both human and mouse (ρ = 0.085 and 0.067 respectively), confirming that more broadly expressed genes evolve under stronger purifying selection. Purifying selection ω_0_ is also negatively correlated to maximum expression, although this is weaker in human, indicating that genes with high expression in at least one tissue have a tendency to evolve under strong purifying selection. More surprisingly, purifying selection ω_0_ is positively correlated to median expression. Note that these are partial correlations; without correcting for other parameters, as expected, ω_0_ correlates negatively with median expression, i.e., highly expressed genes are under strong purifying selection. It appears that this negative correlation is driven by the effect of breadth of expression and of maximum expression, with the residual effect actually in the opposite direction.

**Fig. 1.**
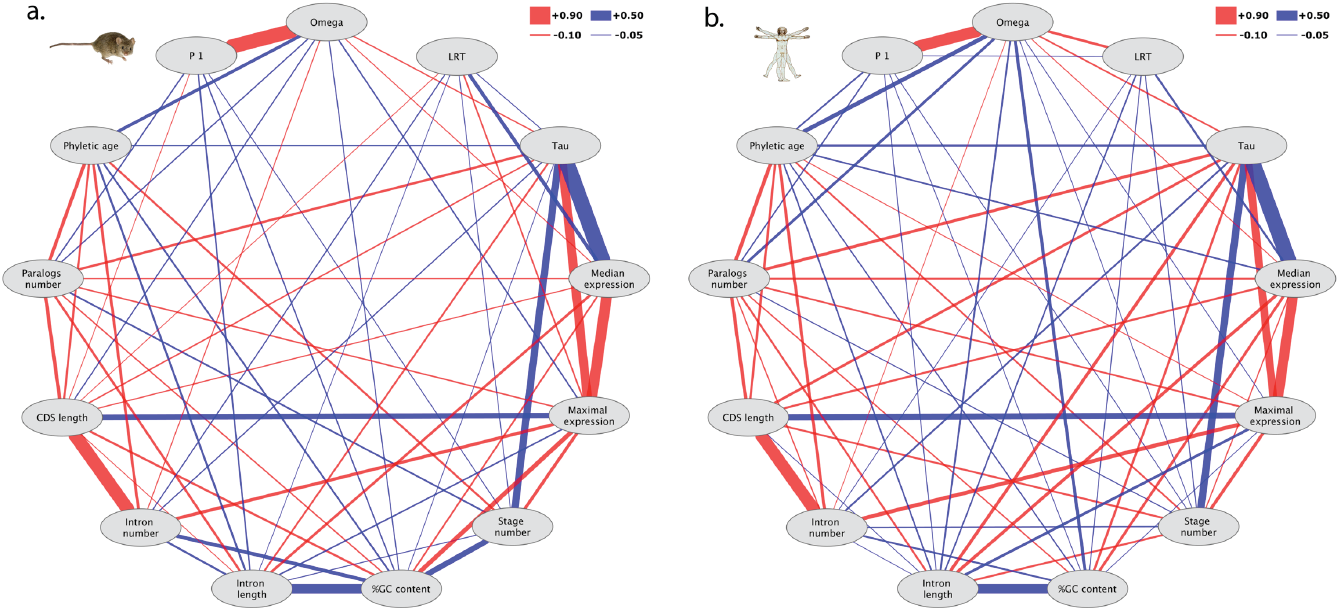

Spearman partial correlations in a) mouse and b) human. The width of the lines shows the strength of correlations. Red lines show positive correlations, blue lines show negative correlations. Only significant correlations (p<0.0005) are shown.

Evolutionary features of the genes, paralog number and phyletic age, have a stronger partial correlation with ω_0_ than expression: older genes, and genes with more paralogs, evolve under stronger purifying selection; again, this is after removing the effect of high levels of expression, as well as the correlation between gene age and number of paralogs. In human, GC content also appears to have a strong influence on ω_0_, but much less so in mouse.

It remains that none of these parameters can explain much of the differences in purifying selection. The total variance of ω_0_ that they explain (using partial Pearson correlation, as Spearman ρ does not relate directly to variance) is 10.2% for human and 13.8% for mouse, thus leaving more than 85% of the variance unexplained.

The strongest correlation with ω_0_ is for p_1_, the proportion of sides evolving neutrally (fig. 1). This partial correlation is ρ = 0.748 in mouse and ρ = 0.598 in human; genes under strong purifying selection have a smaller proportion of sides evolving neutrally. This is not directly due to the way how these parameters are estimated in the branch-site test, since ω_0_ is computed on a distinct set of codons from p_1_. This proportion p_1_ of neutrally evolving sites is otherwise mostly correlated with evolutionary features (phyletic age, paralog number) in human, and with structural features (intron length, GC content) in mouse, but correlations are weak (all ρ < 0.09).

Evidence for positive selection correlates negatively with median expression in both human and mouse (fig. 1), i.e. highly expressed genes are under weaker positive selection (ρ = -0.105 and -0.187 respectively). It should be noted that this correlation concerns relatively weak evidence for selection, since only 4 human and 23 mouse genes in the dataset have significant support for branch-site positive selection (using the false discovery rate of 10% cut-off of Selectome, see Methods).

### Tissue-specific analysis

When the correlation between expression level, selective pressure, and other parameters, is analyzed for each tissue separately, there are large differences, notably in the correlation between expression and purifying selection between tissues (fig. 2).

**Fig. 2.**
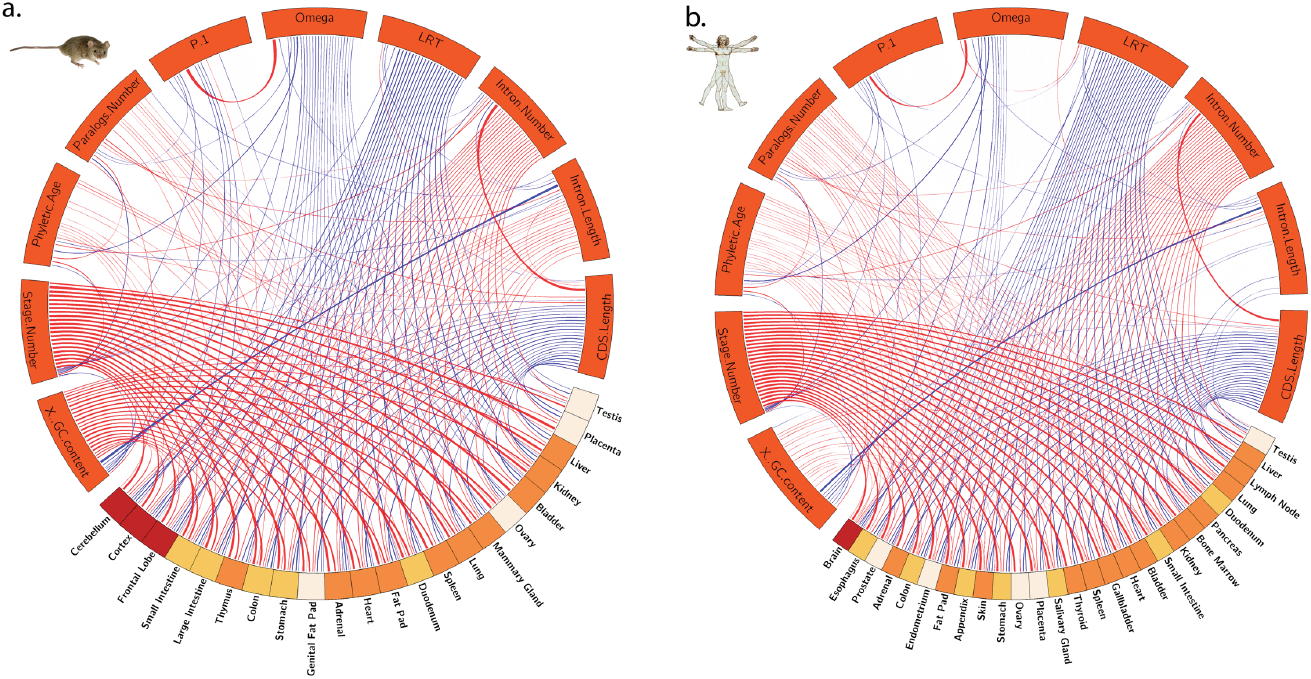

Spearman partial correlation with expression values for each tissue separately for a) mouse and b) human. The width of the lines shows the strength of correlations. Red lines show positive correlations, blue shows negative correlations. Only significant correlations (p<0.0005) are shown. Color of the tissue bands represents different groups of tissues (gastrointestinal system, central nervous system, reproductive system and misc).

In both human and mouse, the strongest correlation with purifying selection ω_0_ is for level of expression in the brain, as expected from previous studies with less tissues (Kuma et al. 1995; Duret & Mouchiroud 2000; Khaitovich et al. 2006; Drummond & Wilke 2008; Tuller et al. 2008). After correcting for all other effects, the residual correlation is rather weak (ρ between -0.065 and -0.107 depending on species and brain part), but always in the direction of stronger purifying selection on genes with higher expression in brain. In human, there are also significant partial correlations for esophagus, prostate, adrenal, colon, and endometrium (fig. 2B). In mouse, there are correlations for all sampled tissues except liver, placenta and testis (fig. 2A); in human the homologous tissues to these three also have among the lowest partial correlations. Interestingly, the only positive partial correlation with ω_0_ is for human testis expression, i.e. higher expression in testis correlates with weaker purifying selection.

The strongest correlations with the proportion of neutrally evolving sites are also for brain tissues, in human and in mouse. Again the correlation is negative, indicating less neutral evolution (i.e., more selection) for more highly expressed genes. There are almost no other tissues with significant partial correlation of expression and p_1_, although for mouse large intestine the correlation is significantly positive.

Concerning evidence of positive selection, on the other hand, there are significant negative partial correlations for all tissues, meaning that for each tissue genes with higher expression have less evidence of positive selection. Brain tissues again have some of the strongest correlations, although they stand out less than for ω_0_ or p_1_. In both mouse and human the correlation is weakest for testis expression, and also quite weak for placenta.

All these correlations include for each tissue both house-keeping and tissue-specific genes; the former might confuse tissue-specific patterns. Thus we repeated the analysis restricted to tissue-specific genes, defined as τ > 0.2 (supplementary fig. S2). The global picture is similar, with notably significant negative partial correlations to ω_0_ only for expression in human brain and mouse cerebellum, significant negative correlation to p_1_ only for mouse brain parts, and conversely significant positive correlations to expression in human and mouse testis.

### Gene age and duplication

As expected, older genes have more paralogs (positive correlation in fig. 1) (Roux & Robinson-Rechavi 2011). Tissue specificity has a rather strong positive partial correlation with paralog number, and a significant weak negative correlation with phyletic age was detected; both correlations are stronger in human than in mouse. That means that, correcting for the correlation between gene age and paralog number, new genes and genes with more paralogs tend to have more tissue-specific expression. While in simple correlation, phyletic age and expression level (median or maximum) have a strong positive correlation (older genes have higher expression), this effect is almost completely lost in the partial correlation, and so is probably spurious.

The phyletic age of the genes correlates negatively with purifying selection but almost no correlation can be seen to neutral evolution or positive selection. This is consistent with previous observations that older genes evolve under stronger purifying selection ((Albà & Castresana 2005, 2007) but see (Elhaik et al. 2006)).

Paralog number correlates negatively with purifying selection in both organisms (-0.064 for mouse and -0.136 for human). This indicates a stronger effect of the biased preservation of duplicates under stronger purifying selection (Brunet et al. 2006; Davis & Petrov 2004; Jordan et al. 2004), than of the effect of faster evolution of duplicated genes (Satake et al. 2012).

### Gene structure

Genes with higher GC content have higher expression level, as shown previously (Urrutia & Hurst 2003), although the effect is not very strong in partial correlation. Previous findings that highly expressed genes are shorter were only partly confirmed: there is a strong negative partial correlation between CDS length and maximal expression, but the partial correlation between median expression and CDS length is weakly positive. Curiously, the partial correlation with intron number is opposite, indicating that genes with high maximum expression tend to have more introns than expected given their CDS length.

### Differences between human and mouse, and between datasets

In general correlations in human are slightly weaker than in mouse, but very consistent (supplementary fig. S3). The strongest difference is between the correlations of GC content and stage number; and of GC content and maximal expression.

There are also noticeable differences between mouse and human in the partial correlations among ω_0_, p_1_ and ΔlnL (evidence for positive selection). In human ΔlnL correlates negatively with p_1_ and positively with ω_0_, indicating that genes with high proportion of neutrally evolving sites and weak purifying selection show little evidence for positive selection. In mouse the correlations are not significant, and in the opposite directions, but the correlation between ω_0_ and p_1_ is much stronger. GC content and paralog number also have stronger correlations to purifying selection in human than in mouse.

We repeated our analyses with large microarray experiments (see Methods, and Supplementary Materials), to control for putative biases in RNA-seq data. There are a few differences, although they do not change our biological conclusions. First, with microarrays tissue-specificity τ appears overall lower, and the correlations between expression parameters (τ, maximal expression, median expression) are stronger. This might be due to the better detection of lowly expressed genes by RNA-seq than by microarray, whereas there seems to be less difference for highly expressed genes (Wang et al. 2014). Conversely, correlations of expression parameters with all other parameters are much stronger for RNA-seq. The correlation between ω_0_ and expression in each tissue separately is stronger with microarrays then with RNA-seq, and significant for all tissues, but the same tissues have the strongest (resp. weakest) correlation between ω_0_ and expression with both techniques. Inversely, the evidence for positive selection has almost no significant correlations with expression in single tissues with microarray data.

We also reproduced our analysis using the precomputed “MI” score for most conserved exon (Rodriguez et al. 2013; Ezkurdia et al. 2014) instead of the branch-site model ω_0_, and all results are similar despite the differences in multiple sequence alignment and in evolutionary model (supplementary fig. S4): e.g., phyletic age is the strongest correlation to MI and median expression has a weak positive partial correlation.

Finally, we repeated our analysis with the RNA-seq data for human and mouse 6 tissues from Brawand et al. (Brawand et al. 2011); results are extremely similar to those with the large RNA-seq experiments used in our main results (Supplementary Material), with less detail of tissues, and less resolution for τ, due to the smaller sampling.

Overall, our results appear quite robust across species and experimental techniques.

## Discussion

### Technical limitations and generality of observations

We use partial correlations to hopefully detect non-spurious relations. Of note, the lack of partial correlation between two parameters does not mean that they are not correlated in practice, but that the correlation is not directly informative, or insufficiently to be detected.

Our analysis was performed on approximately half of the known protein coding genes (9509 for human and 9471 for mouse), for which evolutionary rate could be computed reliably. While this may introduce some bias, it does not appear to have a large influence, since correlations other than to evolutionary rate are very similar on the other half of the coding genes (Supplementary Material).

### Global study of evolutionary rate

Our aim is to understand the causes of variation in evolutionary rates among protein-coding genes in mammals. In yeast or bacteria, the major explanatory feature is the relation between the level of gene expression and purifying selection (Pál et al. 2006; Rocha & Danchin 2004; Rocha 2006). In mammals, firstly levels of expression are more complex to define, due to multicellularity and tissue-specificity, and secondly several other features have been reported to correlate as much or more with evolutionary rate, in studies which did not necessarily incorporate all alternative explanations.

In this study, we have focused on the dN/dS ratio, or ω, and distinguished further the three forces which affect this ratio, under the classical assumption that dS is overwhelmingly neutral (although see (Rubinstein et al. 2011; Macossay-Castillo et al. 2014; Dimitrieva & Anisimova 2014)). The intensity of purifying selection is clearly the main component of the overall ω: on average more than 85% of codons are in the purifying selection class of the evolutionary model used. Analyzing separately neutral evolution and purifying and positive selection, we find that (i) these three forces do not affect protein coding genes independently, and (ii) they have different relations to gene expression and to other features. Notably, genes which are under stronger purifying selection have less codons predicted under neutral evolution. Importantly, we computed evolutionary rates on filtered alignments (Moretti et al. 2014), which probably eliminates mostly neutrally evolving sites, thus underestimating p_1_. Still, it appears that to the best of our knowledge these two forces act in the same direction. The relation is less clear concerning the evidence for positive selection, with opposite correlations in human and mouse. But we are limited by the weak evidence for positive selection on the branches tested, at the human-chimpanzee and mouse-rat divergences. Overall, these relations between forces acting on ω_0_ deserve further investigation with more elaborate evolutionary models (e.g., Murrell et al. 2012; Zaheri et al. 2014). Despite the limitations of the estimation of positive selection, this is the component of evolutionary rate which has the strongest partial correlation with the level of gene expression, both with the median expression over all tissues, and with expression in brain tissues. This implies that when expression patterns constrain the protein sequence, they also strongly limit adaptation (strong purifying selection and very low positive selection).

So what explains evolutionary rate? The strongest partial correlation of ω_0_ is with phyletic age: older genes evolve under stronger purifying selection. While the use of partial correlation allows us to correct for some obvious biases in detecting distant orthologs, such as gene length, we cannot exclude that results be partially caused by the easier detection of orthologs in distant species for proteins with more conserved sequences (Elhaik et al. 2006; Albà & Castresana 2007; Moyers & Zhang 2014). I.e., genes with weak purifying selection may be reported as younger than they are, because the orthologs were not detected by sequence similarity. We obtain similar results with an exon-based index of sequence conservation, MI (supplementary fig. S4). Whatever the contributions of methodological bias and biological effect, this correlation is not very informative about causality, since stronger selection will not be caused by the age of the gene.

The next strongest partial correlation with ω_0_ is the GC content of the gene. In mammals, the variation in GC content of genes seems mostly due to GC-biased gene recombination (Montoya-Burgos et al. 2003), and this in turn has been show to impact estimation of dN/dS (Galtier et al. 2009). But while GC-biased gene recombination is expected to lead to high GC and an overestimation of ω, we find a negative correlation between ω_0_ and GC content, consistent with previous observations in Primates (Bullaughey et al. 2008). Of note, estimating the actual biased recombination rate rather than GC content is limited by the rapid turn-over of recombination hotspots (Glémin et al. 2014), and recombination rate appears to have only a very weak effect on dN/dS in Primates once GC content is taken into account (Bullaughey et al. 2008). This could be seen in our study, as recombination rate did not show any significant correlation to any of the parameters, except GC content (Supplementary Material). The previously reported relation between dN/dS and intron length seems to be mostly an indirect effect of the strong correlation between GC content and intron length (Montoya-Burgos et al. 2003; Duret et al. 1995).

The significant, although weaker, partial correlation of ω_0_ to paralog number is consistent with previous observations that genes under stronger purifying selection are more kept in duplicate (Davis & Petrov 2004; Yang & Gaut 2011; Jordan et al. 2004; Brunet et al. 2006). The level of gene expression has been reported repeatedly to be the main explanatory variable for dN/dS (Subramanian & Kumar 2004; Liao & Zhang 2006; Pál et al. 2001; Drummond et al. 2005; Wall et al. 2005), notably in *S. cerevisiae*. Our first observation is that no aspect of expression in human and mouse adult tissues is as strong an explanatory factor for any component of evolutionary rate as what was reported in yeast. Our second observation is that three aspects of expression influence evolutionary rate most strongly: breadth of expression τ; number of developmental stages (fig. 1; table 1); and expression in brain tissues (fig. 2). The third, surprising, observation is that median expression is positively correlated with ω_0_: taking into account other parameters, genes which have higher expression on average are under weaker purifying selection; whereas the correlation with maximal expression is negative, as expected. Thus in mammals the negative correlation between median expression and evolutionary rate appears to be an indirect effect of stronger selection on broadly expressed genes and on genes with high maximal expression in at least one tissue (this is also true if we take the mean instead of median expression, see Supplementary Materials).

We confirm previously reported observations that expression breadth is more important then expression level itself in mammals (Park & Choi 2010). dN was previously found to be threefold lower in ubiquitous than in tissue-specific genes, while dS did not vary with expression specificity (Duret & Mouchiroud 2000). Other studies indicate that genes expressed in few tissues evolve faster then genes expressed in a wide range of tissues (Liao & Zhang 2006; Gu & Su 2007; Park & Choi 2010), or that tissue-specific genes have more evidence for positive selection (Haygood et al. 2010). In mouse, but not human, τ is weakly negatively correlated to evidence for positive selection: broadly expressed genes seem to be more affected by positive selection, *contra* Haygood et al (Haygood et al. 2010). We also notice that tissue-specificity and maximal expression are correlated, i.e. more tissue-specific genes have higher maximum expression in one tissue. Thus these two forces appear to act on different genes: some genes are under strong purifying selection because they are broadly expressed, suggesting an important role of pleiotropy, while other genes are under strong purifying selection because they are highly expressed in few tissues, suggesting an important role of the tissue-specific optimization of protein sequences. Of note, analyzing separately only brain expression relative to maximal and breadth of expression in other tissues gave similar results, thus brain expression alone is not driving these patterns (not shown).

Some studies have reported that expression level and tissue specificity are less important than gene compactness and essentiality in mammals (Liao et al. 2006). Liao et al. (Liao et al. 2006) reported that compact genes evolve faster, but this correlation is very weak in our study. We could not either confirm that highly expressed genes are shorter (Li et al. 2007; Chen et al. 2005; Urrutia & Hurst 2003). We have used the longest transcript for each protein coding gene, as evolutionary parameters (ΔlnL, ω_0_, p_1_) were calculated for the transcript. But this might not be the transcript most expressed and used in all tissues (Gonzalez-Porta et al. 2013). We repeated calculations with the most expressed transcript (Supplementary Material), but results were unchanged; we show these results only in supplementary materials, as the estimation of transcript-level expression does not yet appear to be very reliable (Lahens et al. 2014; Cho et al. 2014). Finally, we tried to investigate the impact of essentiality, but we found no significant effect (supplementary fig. S5); we note that we have very low power to test this effect, especially in human.

The analysis was also performed adding connectivity and recombination data, for mouse only. This reduced the number of analyzed genes to 4599 (Supplementary Material). Correlations were mostly unchanged, with the largest difference being for the correlation between stage number and phyletic age, from 0.11 to 0.093. No notable change of correlations with ω_0_ was detected, and connectivity and recombination rate do not show any significant correlation to evolutionary rate.

The largest partial correlations that we observed for components of evolutionary rate are between brain expression level and evidence for positive selection, at -0.203 to -0.188 in mouse (for different brain parts), and -0.168 in human (whole brain). For purifying selection we find weaker but significant partial correlations with brain expression and with the number of stages, between 0.065 and 0.119. And brain tissues also have the strongest partial correlation over expression in tissues for neutral evolution (fig. 2). It has been previously reported that brain expression is a major component of evolutionary rate in mammals and other animals (Khaitovich et al. 2006; Duret & Mouchiroud 2000; Kuma et al. 1995; Drummond & Wilke 2008), and here we confirm the dominance of this component, even taking other effects into account. Importantly we show that this affects all forces acting on protein evolutionary rate: purifying selection, neutral evolution, and positive selection. Thus the median expression of genes over more than 20 tissues is a poor explanation of protein evolutionary rate, relative to brain expression.

### Tissue specific patterns

There are striking differences between tissues in the extent of the correlations with structural and evolutionary parameters. As already mentioned, brain tissues present the strongest partial correlations with evolutionary rate; results are consistent when only tissue-specific genes are used. We observe this for the three evolutionary forces estimated. In most comparisons, the correlation is stronger for brain expression than for any global measure of expression. This is consistent with the translational robustness hypothesis, which proposes that highly expressed genes are under stronger pressure to avoid misfolding caused by translational errors, thus these genes are more conserved in evolution (Drummond et al. 2005), and that neural tissues are the most sensitive to protein misfolding (Drummond & Wilke 2008). This slow evolution of genes expressed in neural tissues has been repeatedly reported (Duret & Mouchiroud 2000; Kuma et al. 1995; Necsulea & Kaessmann 2014), especially for the brain (Park & Choi 2010); it has also been related to higher complexity of biochemical networks in the brain than in other tissues (Kuma et al. 1995).

Fast evolution of genes expressed in testis is also well documented (Khaitovich et al. 2006; Brawand et al. 2011; Necsulea & Kaessmann 2014), and could be due to lower purifying selection, an excess of young genes and leaky expression, or to positive selection due to sexual conflict. We observe neither a stronger correlation between expression in sexual tissues and evidence for positive selection, nor a stronger correlation between expression in sexual tissues and the proportion of sites evolving neutrally. What we do observe is that the weakest partial correlation between expression in a tissue and purifying selection is for testis, and that it is also quite weak for placenta, with even a surprising positive correlation between ω_0_ and expression in human testis, which remains when only tissue-specific genes are used. This is consistent with the “leaky expression” model: being expressed in the testis does not appear to be an indicator of function carried by the protein sequence. Interestingly, expression in testis is negatively correlated with the number of paralogs, significantly so in mouse: genes which are more expressed in testis have less paralogs, after correcting for other effects.

While the strong correlation of ω_0_ with expression in the brain, and the weak correlation with expression in testis are expected, we also observe less expected patterns. Most notably, liver expression has the next weakest correlation with ω_0_ after testis (and placenta in mouse). Although it was reported before that liver expressed genes are evolving faster (Khaitovich et al. 2006; Duret & Mouchiroud 2000), it was reported with much fewer tissues, and not highlighted. Liver expression is also positively correlated with the proportion of neutral sites, unlike brain or testis expression, although this is not significant. Interestingly, liver has the strongest correlation of expression with phyletic age, implying that despite low purifying selection, old genes are more expressed in liver. In any case, this outlier position of liver has important practical implications, since liver is often used as a “typical” tissue in studies of gene expression for molecular evolution (e.g., (Gilad et al. 2006; Blekhman et al. 2010; Enard et al. 2002)).

## Conclusion

The main result of our study is that average adult gene expression is quite lowly informative about protein evolutionary rate, while purifying selection on genes highly expressed in the brain and breadth of expression are our best bets for a causal factor explaining evolutionary rates. A practical consequence is that great care should be taken before using expression from other tissues, including widely used ones such as liver, as proxies for the functional importance of mammalian genes.

Finally, all calculations were performed with expression in adult tissues. It is possible that expression in embryonic development be more important for evolutionary constraints in mammals, and this should be explored further.

## Supplementary Material

The most important Supplementary Materials are available at Genome Biology and Evolution online (http://gbe.oxfordjournals.org/). Data sets and other supplementary figures are available at: http://dx.doi.org/10.6084/m9.figshare.1221771.

## Acknowledgments

We thank Julien Roux for helpful comments on the manuscript. This work was supported by the Swiss National Science Foundation (grants number 31003A 133011/1 and 31003A_153341/1) and Etat de Vaud. The computations were performed at the Vital-IT Center (http://www.vital-it.ch) for high-performance computing of the SIB Swiss Institute of Bioinformatics.

